# RADIAL AND TANGENTIAL FIBER COMPONENTS IN HUMAN SUPERFICIAL WHITE MATTER AND THEIR ASSOCIATIONS WITH MACROSCALE FUNCTION

**DOI:** 10.64898/2026.10.02.756249

**Authors:** Youngeun Hwang, Laura Vavassori, Paul Bautin, Raul Rodriguez-Cruces, Edith Gaspar-Martinez, Donna Gift Cabalo, Alexander Ngo, Meaghan Smith, Robert Leech, Ricardo Coronado-Leija, Laurent Petit, Paolo Avesani, Silvio Sarubbo, Alan C Evans, Luis Concha, Boris C. Bernhardt

**Author notes:** Corresponding authors: Youngeun Hwang, MSc Boris C. Bernhardt, PhD.

## Abstract

The superficial white matter (SWM), a thin layer of white matter beneath the cortex, contains a complex arrangement of short- and long-range fiber systems that support communication between cortical regions. These fiber populations include subcortical U-shaped fibers (U-fibers) that predominantly follow a tangential trajectory along the gray matter-white matter interface and long-range fibers that extend more radially through the deeper white matter to connect distant cortical regions. Here, we developed a surface-based framework to dissociate radial and tangential fiber components within the human SWM and investigated their microstructural properties and functional relevance using *in-vivo* high-field MRI. These components were quantified using apparent fiber density (AFD), fractional anisotropy (FA), and mean diffusivity (MD). We further contextualized SWM fiber architecture by examining depth-dependent microstructural profiles and their relationship with normative patterns of cortical organization and hierarchical specialization. The anatomical assessment of the proposed fiber orientation separation was supported by tractography-based bundle analyses, while the anatomical accuracy of tractography reconstructions was confirmed through comparison with *post-mortem* white matter dissection data. To assess their functional relevance, the fiber predominance map derived from radial and tangential AFD was related to spatial patterns of functional variability across the cortex. Regions with greater radially oriented fiber contribution exhibited more gradual changes in functional similarity across geodesic distance, whereas tangentially oriented fiber-enriched regions showed more rapid local functional differentiation. These findings reveal distinct structural and functional signatures of radial and tangential SWM fiber components and provide a framework for investigating their contributions to cortical organization and functional specialization.

**SIGNIFICANCE STATEMENT:** The human cerebral cortex is organized into specialized regions that support diverse functions, but the structural principles underlying this organization remain incompletely understood. The superficial white matter (SWM), a thin layer of white matter beneath the cortex, contains intermingled short- and long-range fiber systems that support communication between cortical regions. However, these fiber populations have remained difficult to characterize because they are closely intertwined within the SWM. By developing a reproducible framework for mapping and quantifying distinct SWM fiber systems, this work provides a foundation for systematic investigation of this complex compartment. Integration of multimodal datasets, together with open sharing of analytical tools and standardized features, promotes transparency, reproducibility, enabling future studies of how superficial connectivity supports brain organization.

## INTRODUCTION

The superficial white matter (SWM), a thin layer of white matter beneath the cortex, is a structurally complex compartment that contains diverse fiber populations supporting communication between cortical regions. These pathways can be broadly characterized according to their spatial extent and orientation within the SWM, including short-range fibers that course tangentially along the gray matter-white matter (GM/WM) interface and longer-range fibers that extend more radially through the deeper white matter to connect spatially separated cortical regions (Van Dyken et al., 2024; Garcia-Saldivar et al., 2024). Specifically, subcortical U-shaped fibers (U-fibers) interconnect adjacent cortical gyri, forming dense short-range connections between them (Movahedian et al., 2020; Shastin et al., 2022; Schüz et al., 2002; Kirilina et al., 2020; Gahm et al., 2019). In contrast, long-range association fibers extend beyond neighboring gyri and contribute to communication between more distant cortical regions through deep white matter pathways (Schüz et al., 2002; Schmahmann et al., 2009). Together, these radially and tangentially oriented fiber populations form an intricate architecture within the SWM that supports both local cortical interactions and large-scale cortical communication.

Differences in the organization and developmental trajectories of these fiber populations may contribute to their distinct roles across the lifespan. U-fibers primarily arise from pyramidal neurons in the superficial cortical layers and develop in parallel with cortical gyrification and sulcation during the second half of gestation (Yoshino et al., 2020; Schüz et al., 2002; Schmahmann et al., 2009). Unlike many long-range fibers, they undergo prolonged maturation and remain incompletely myelinated well into later stages of life (Movahedian et al., 2020), suggesting continued involvement in maturation, experience-dependent plasticity, and aging processes (Parazzini et al., 2002). Beyond their significance for these development and maturation, which may be further highlighted by differences in U-fiber organization across species, SWM U-fibers have important clinical implications. U-fiber configurations have been shown to differentiate white matter pathologies (Riley et al., 2018), as the extent and pattern of involvement of these superficial fiber systems may vary across white matter disease processes. More broadly, alterations in SWM organization and microstructure, as well as U-fiber density, have been associated with neurological and neurodevelopmental conditions, including epilepsy (Larivière et al., 2020; O’Halloran et al., 2017; Liu et al., 2016), autism spectrum disorder (Hong et al., 2019; Hau et al., 2019, 2022; Zikopoulos et al., 2010), and Alzheimer’s disease (Carmeli et al., 2014). These findings highlight the potential importance of local U-fiber pathways, in addition to long-range fibers, in maintaining normal cortical organization and in contributing to network dysfunction. However, quantitative characterization of U-fibers and their relationship to long-range fiber systems remains limited (Guevara et al., 2017, 2020).

A major technical challenge in characterizing SWM organization is the close intermingling of radially and tangentially oriented fiber populations within this anatomically complex compartment (Hwang et al., 2026; Kirilina et al., 2020). *Post-mortem* dissection studies have provided valuable direct insights into the anatomical organization of white matter fiber bundles, however, the availability of high-resolution three-dimensional datasets suitable for systematic and quantitative analyses remains limited. Consequently, *in-vivo* neuroimaging approaches have become essential for characterizing SWM organization across individuals. Recent advances in magnetic resonance imaging (MRI), have enabled increasingly precise and non-invasive investigation of brain microstructure, including the thin SWM layer (Hwang et al., 2026). Diffusion-weighted MRI (dMRI), in particular, provides a means of characterizing the orientation and structural connectivity of white matter pathways (Maier-Hein et al., 2017; Le Bihan and Breton, 1985; Jeurissen et al., 2019). Importantly, developments in high-angular resolution dMRI have improved the ability to resolve multiple fiber populations within crossing-fiber regions, providing microstructural information beyond the limitations of diffusion tensor imaging (Maier-Hein et al., 2017; Frank, 2001; Tuch et al., 2002; Tournier et al., 2004, 2007, 2019; Jeurissen et al., 2014; Wu et al., 2020; Garyfallidis et al., 2014). The multimodal nature of MRI further enables integration of structural and functional imaging, creating opportunities to examine how distinct SWM fiber components relate to cortical functional organization in the living human brain.

The study of task-free, or resting-state, functional MRI (fMRI) permits the identification of intrinsic functional networks, enabling investigation of how local and large-scale brain function relates to underlying white matter architecture (Biswal et al., 1995; Raichle, 2015; Smith et al., 2009; Yeo et al., 2011; Cauda et al., 2011; Margulies et al., 2016). Consistent with this perspective, lesion-based studies have reported that alterations affecting the SWM and U-fiber systems are associated with atypical functional connectivity patterns (Shukla et al., 2011; Liu et al., 2016; Hong et al., 2019), highlighting the potential contribution of the SWM compartment to large-scale brain organization. Increasing research suggests that functional organization is structured along continuous spatial gradients that reflect hierarchical and distributed aspects of brain organization (Margulies et al., 2016; Paquola et al., 2019; Smallwood et al., 2021; Burt et al., 2018; Sydnor et al., 2023). More recently, spatial statistical methods have characterized distance-dependent patterns of functional similarity across the cortex, providing insight into how brain geometry and structural organization constrain the spatial patterning of brain function (Leech et al., 2023; Spence et al., 2007; Bowman, 2007; Worsley et al., 1998). Utilizing these approaches, it is now possible to comprehensively investigate how SWM microstructural architecture relates to functional specialization and integration across the cortex (Hwang et al., 2026). This provides an opportunity to better understand the interplay between SWM properties and cortical functional organization.

In this study, we developed and assessed a novel framework for distinguishing radial and tangential fiber components within the human SWM, providing an *in-vivo* approach to characterize putative long-range and U-fiber systems, respectively. Specifically, we operationalized an open-source pipeline that uses Laplacian streamlines as a local reference for fiber orientation, enabling the systematic separation of radial and tangential components across SWM depths in human MRI datasets. Using a surface-based approach, this framework provides consistent depth-resolved sampling while accounting for the complex folding geometry of the cortex across subjects. We quantified the microstructural properties of these components using apparent fiber density (AFD), fractional anisotropy (FA), and mean diffusivity (MD), and examined their depth-dependent profiles across the SWM. To further contextualize these diffusion-derived profiles, we assessed their relationship with normative patterns of cortical organization and functional specialization, as defined by established anatomical classifications. Next, we evaluated this framework using tractography-derived bundle reconstructions and compared them against *post-mortem* brain dissection photogrammetry data to evaluate the biological plausibility of the proposed separation approach within the SWM. Finally, we tested whether SWM fiber architecture was associated with cortical functional organization by examining its relationship with spatial patterns of functional connectivity. To facilitate reproducibility and independent use of the framework, we publicly release all analysis code together with standardized SWM feature maps generated in this study.

## RESULTS

### Distinct organization of radial and tangential fiber systems across SWM depths

We introduce an open-access framework to disambiguate the contributions of radial and tangential fiber components within the human SWM. The framework utilizes streamlines extending from the GM/WM interface toward the ventricular walls, which follow the Laplacian field across the white matter. These streamlines, hereafter referred to as Laplacian streamlines, provide a local orientation reference within the SWM against which dMRI-derived fiber orientations can be compared. Specifically, fiber orientation distribution function (fODF) lobes, representing distinct fiber orientations within a voxel, were classified into radial and tangential components according to their orientation relative to the Laplacian streamlines (**Fig. 1B**). Components oriented parallel to the Laplacian streamlines were interpreted as radial fibers extending into the deeper white matter to connect spatially separated cortical regions through afferent and efferent projections, whereas perpendicular components were considered tangential to the GM/WM interface. These radial and tangential components were subsequently quantified using AFD, defined as the integral of the corresponding fODF lobe, across SWM depths, extending from the GM/WM interface to 3 mm into the white matter (**Fig. 2**). FA and MD were also independently estimated for the radial and tangential components using the multi-resolution discrete search (MRDS) approach, which enables diffusion metrics to be resolved along distinct fiber orientations (Coronado-Leija et al., 2017).

**Figure 1.**
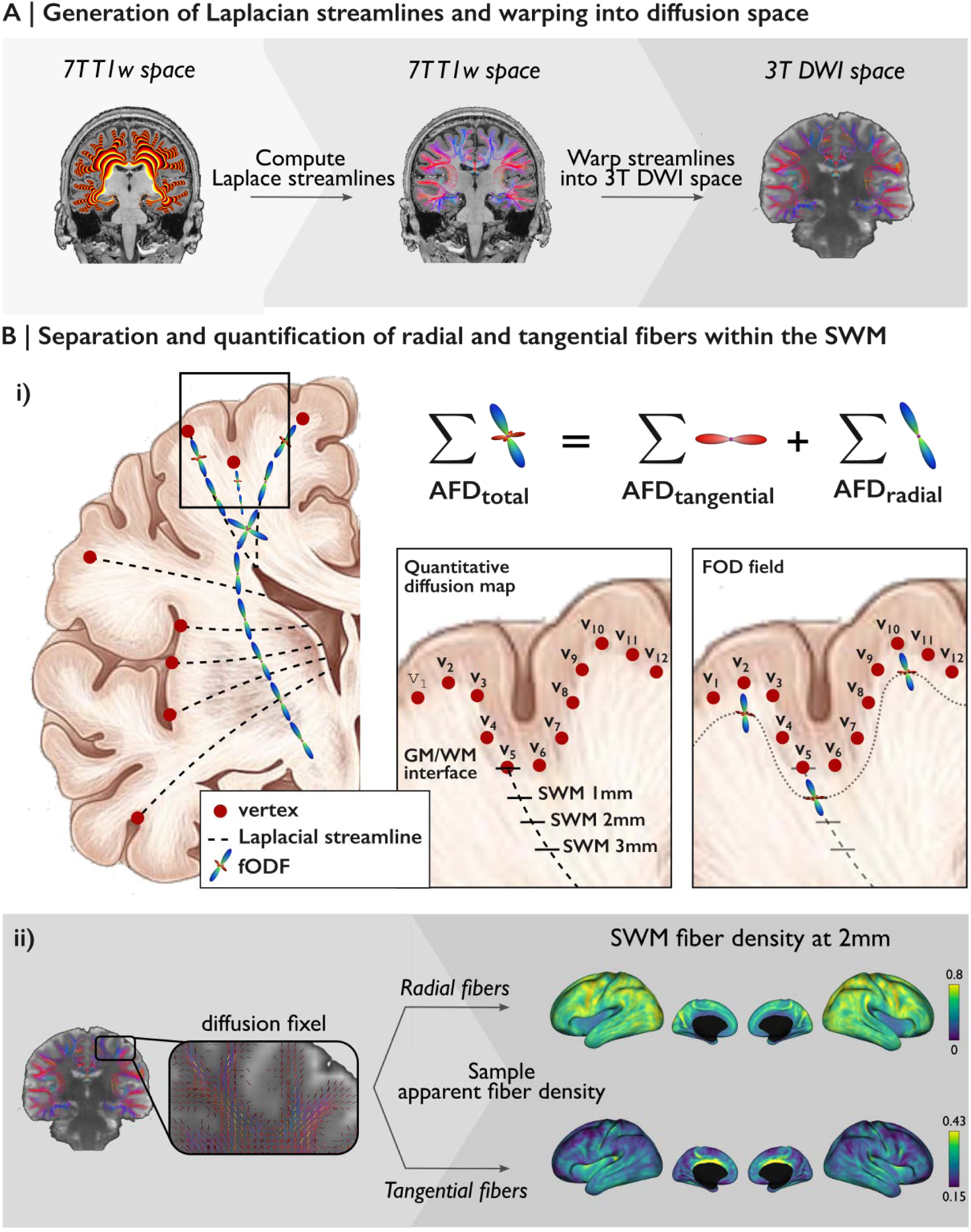
Surface-based framework for SWM fiber separation and quantification. **(A)** The Laplacian field within the white matter domain was solved using the “*aparc+aseg*” segmentation derived from each 7T T1w to sample Laplacian streamlines at each cortical vertex. The resulting streamlines were subsequently warped into 3T DWI space. **(B) *(i)*** We assumed that U-fibers predominantly follow the GM/WM interface in a tangential orientation, whereas long-range fibers approach the cortex in a more radial orientation in the SWM. Per-bundle apparent fiber density (AFD) is derived from the integral of corresponding fiber orientation distribution function (fODF) lobes. ***(ii)*** Diffusion fODFs and resulting AFD were separated based on their alignment with the Laplacian streamline directio : fODF lobes with the largest dot product with the streamline and angular deviations of less than 40° at gyral vertices or 50° at sulcal vertices were classified as the radial component, while the remaining fODF lobes were averaged t represent the tangential component. Separation of multiple tensors and their resulting metrics was performed in the same fashion (described in **Methods** section). Panel **B*(i)*** adapted from Garcia-Saldivar et al., 2024.

**Figure 2.**
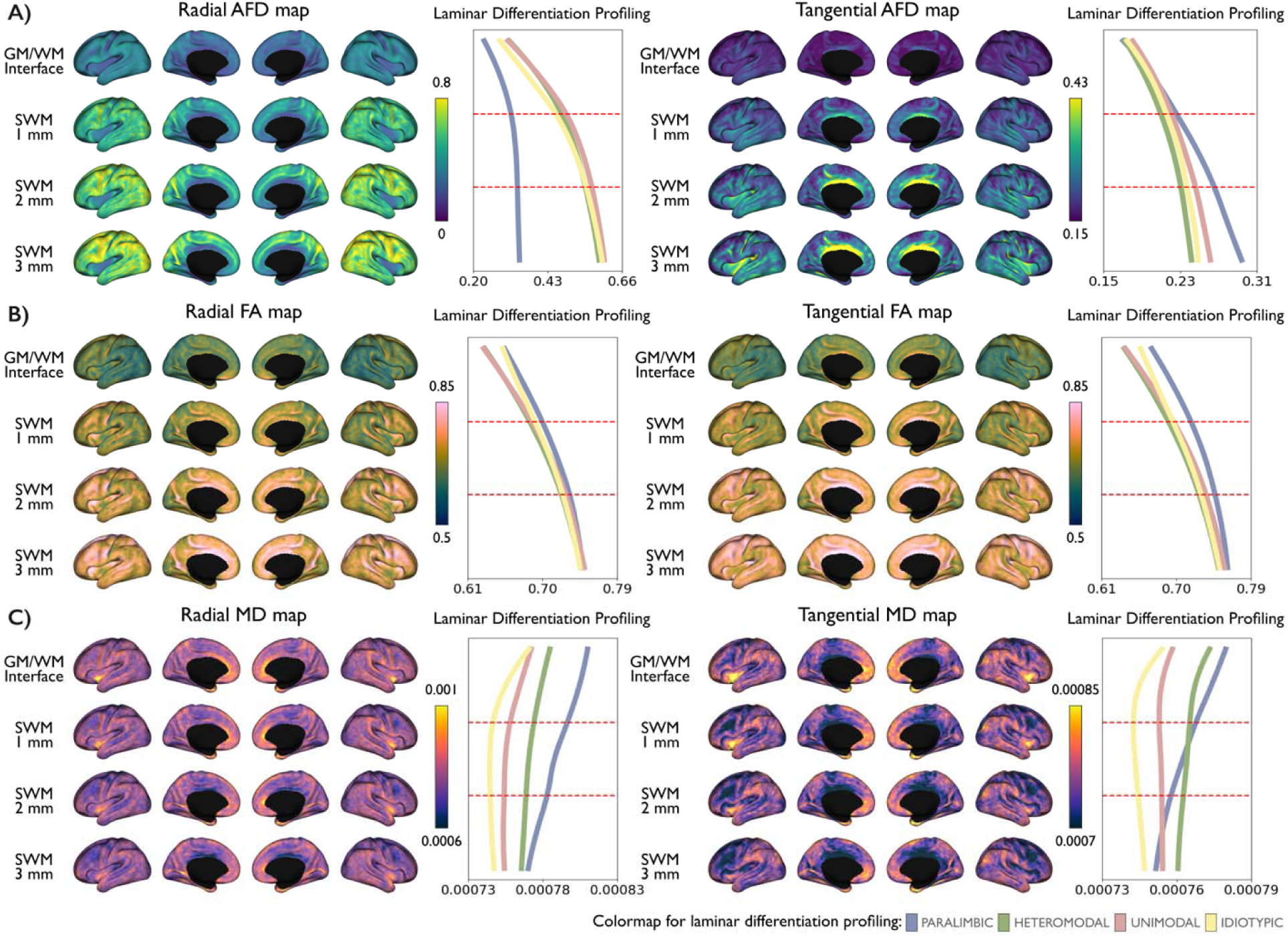
Quantitative diffusion metrics of radial and tangential fiber components within the SWM. In addition to dissociated radial and tangential **(A)** AFD maps, **(B)** FA and **(C)** MD maps were estimated for each component using the MRDS framework to characterize fiber-specific diffusion properties within each voxel. Radial fiber component is interpreted as afferent and efferent projections extending into the deeper white matter to connect spatially distant cortical regions (i.e., long-range fibers), whereas the tangential component is consistent wit putative U-fibers. Spatial relationships between diffusion-derived intensity profiles and laminar differentiation are also presented. Red dotted lines indicated every 1 mm within the SWM.

As shown in **Fig. 2A**, qualitative examination of laminar differentiation revealed notable density differences in paralimbic regions compared with other cortical classes, for both radial and tangential components. In these regions, radial AFD values were relatively lower compared with other cortical areas, whereas tangential AFD exhibited slightly higher intensity. Similar findings were observed in the available 7T diffusion MRI data (**Supplementary Fig. 1**). FA maps (**Fig. 2B**) showed similar patterns for both components, with FA values increasing progressively with SWM depth across brain regions. In contrast, MD maps (**Fig. 2C**) demonstrated clear regional and depth-dependent variations. Notably, MD values in paralimbic regions exhibited a pronounced decrease into the white matter. In this analysis, subject-level variability remained low across cortical classes (**Supplementary Table 1**). Together, these findings reveal a distinct depth-dependent SWM microstructural profile in paralimbic regions. This organization parallels the transitional cortical architecture of paralimbic regions described in classical hierarchical models (Mesulam et al., 1998). The accompanying depth-dependent reduction in MD may further reflect increased microstructural complexity within paralimbic SWM, consistent with previous observations based on quantitative MRI contrast mapping (Hwang et al., 2026).

### Anatomical assessment of fiber tracts and their orientations relative to Laplacian streamlines

We measured the angles between tractography-derived fiber bundles and the corresponding Laplacian streamlines to characterize orientation differences between radially and tangentially oriented fibers in the SWM. We focused on the connectivity between the precentral gyrus (PrCG) and postcentral gyrus (PoCG) as a representative tangentially oriented short-range fiber bundle, and the connectivity between the middle frontal gyrus (MFG) and angular gyrus (AG) as a representative radially oriented long-range fiber bundle. Consistent with the proposed framework, tangentially oriented fibers exhibited angles greater than 50° in sulcal regions and 40° in gyral regions relative to the Laplacian streamlines within th first 2 mm of the SWM (0-1 mm: 56.11 1.83 and 1-2 mm: 49.31 1.76 at the whole-brain level). In contrast, radially oriented fibers showed angles smaller than 50° in sulcal regions and 40° in gyral regions over the same depth range (0-1 mm: 43.57 2.33 and 1-2 mm: 41.66 3.05 at the whole brain level). (**Fig. 3A**; see **Supplementary Table 2** for results at the gyral and sulcal vertices).

**Figure 3.**
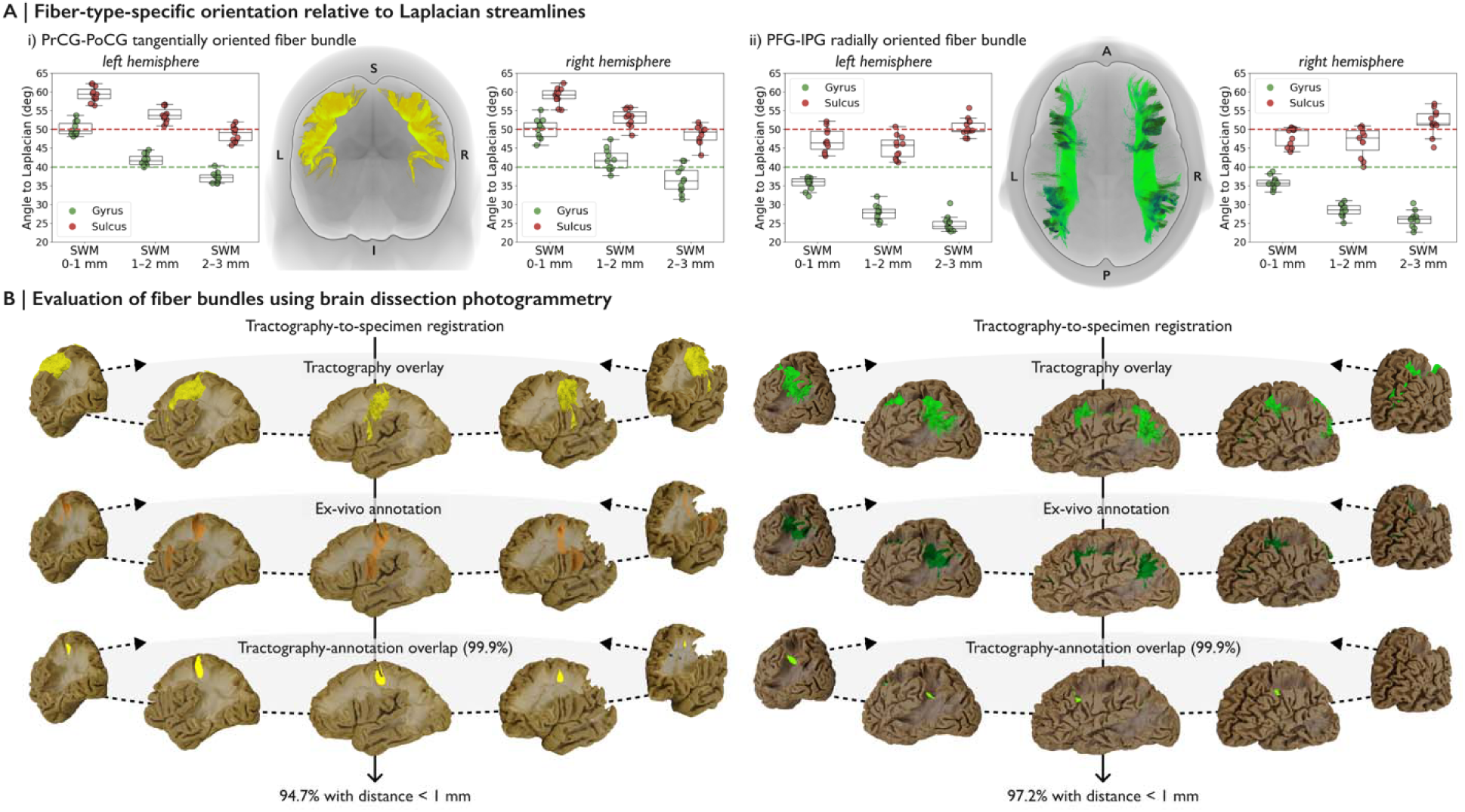
Anatomical validation of the surface-based framework for SWM fiber separation using *post-mortem* anatomically confirmed *in-vivo* tractography. (A) To evaluate the orientation-based separation framework in the SWM, the angles between Laplacian streamlines and two representative fiber pathways for each radial and tangential fiber component were calculated within the first 3 mm of the white matter. (B) Representative *in-vivo* fiber tracts were anatomically validated using brain dissection photogrammetry by registering reconstructed bundles to each specimen space and measuring the spatial distance between tractography-derived pathways and manually annotated white matter pathways from the *post-mortem* dissection data. Black dotted lines indicate the direction of the change in viewing angle.

To further assess the anatomical validity of these fiber tracts, we compared tractography reconstructions with *post-mortem* dissection photogrammetry data from the *BraDiPho* repository (https://bradipho.eu). Following an established anatomical evaluation pipeline for *in-vivo* tractography (Vavassori et al., 2025a), reconstructed fiber tracts were transformed into the space of each ten dissection specimens and visually and quantitatively compared with manual annotations of the corresponding white matter connections. Qualitative evaluation first examined whether the tractography reconstructions encompassed the anatomical extent of the pathways identified in the dissections. Subsequently, quantitative agreement between the two modalities was assessed by calculating the proportion of annotation points located within 1 mm of the tractography-defined overlap region. The spatial distance between reconstructed tracts and annotations was computed for each specimen and across different dissection layers (**Fig. 3B**; se **Supplementary Fig. 2** for full distributions).

Overall, both *in-vivo* radially and tangentially oriented fiber bundles showed high spatial correspondence with the annotations derived from *post-mortem* dissection data, supporting the anatomical reliability of the tractography reconstructions. In addition, orientation analysis relative to the Laplacian streamlines demonstrated a clear distinction between the two fiber orientation systems. Tangentially oriented fibers predominantly showed angles above 40° in gyral regions and 50° in sulcal regions, whereas radially oriented fibers showed angles below these respective thresholds.

### Relationship between SWM fiber organization and cortical functional transitions

We examined whether SWM fiber architecture was associated with cortical spatial dependencies in functional organization. Specifically, we assessed the relationship between functional variability and spatial distance across the cortex to determine whether spatial autocorrelation properties reflected regional differences in radial and tangential fiber predominance within the SWM (**Fig. 4**).

**Figure 4.**
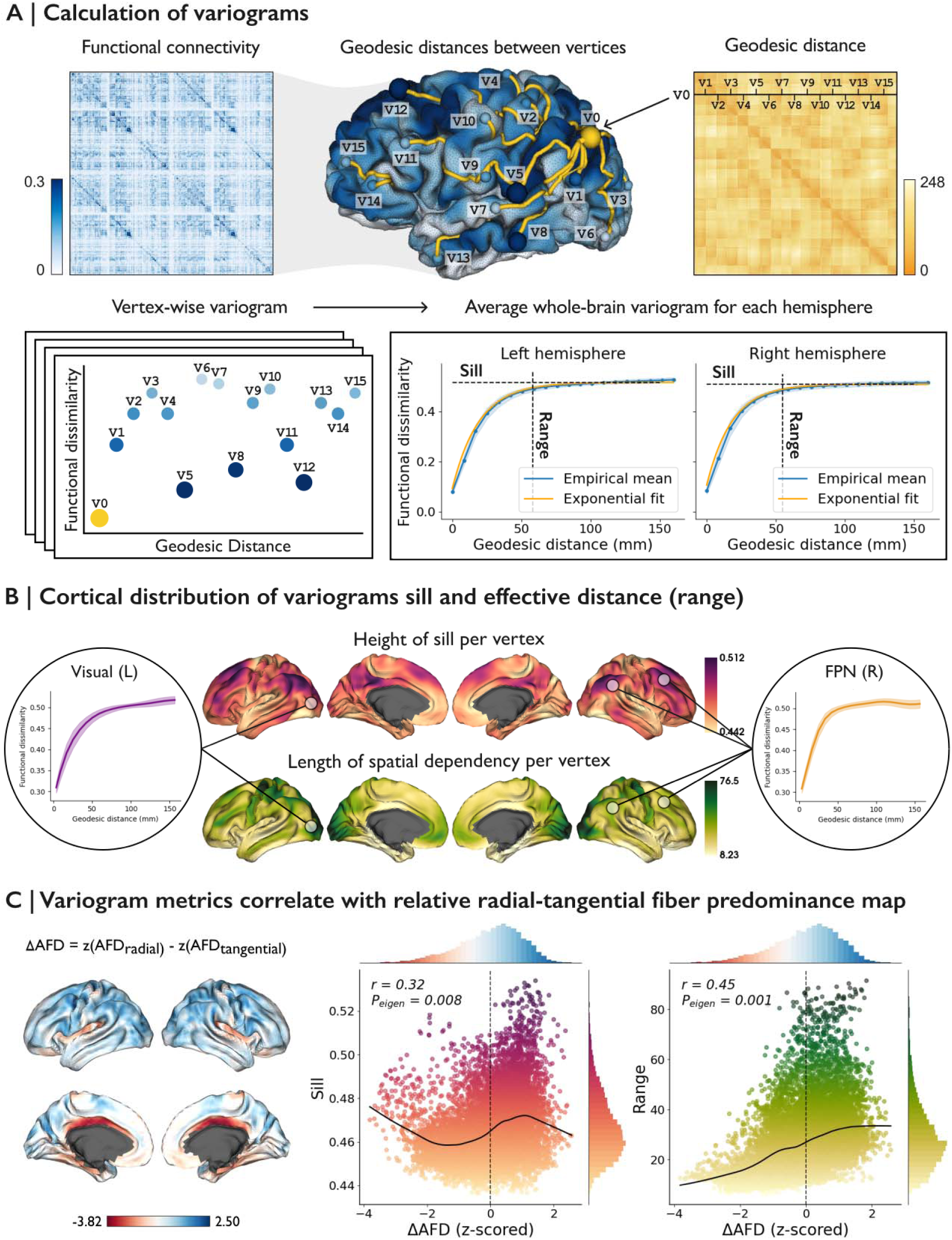
Functional relevance of SWM microstructure and cortical functional variability. **(A)** Schematic illustration of spatial variograms used to characterize how functional connectivity changes as a function of geodesic distance between cortical regions. Representative whole-brain variograms are shown in the *bottom right*, with the sill indicating the asymptotic correlation between vertices and the range representing the geodesic distance (mm) at which the asymptote is reached. **(B)** Mean vertex-wise sill and range maps averaged across subjects. **(C)** The radial-tangential fiber predominance map shown in the *left* panel was generated by subtracting z-scored tangential AFD maps from radial AFD maps at 2 mm SWM depth. The *right* panel shows correlations between variogram-derived metrics and fiber predominance index. **Abbreviations:**, *r*, Spearman correlation;, *P_eigen_* significance.

Spatial variogram analysis was performed following a previously established framework (Leech et al., 2023; Vos de Wael et al., 2020; Burt et al., 2020). Variograms provide a compact representation of how functional organization varies across anatomical distances, capturing both the distance over which spatial dependencies persist and the magnitude of spatial variation. Empirical vertex-wise variograms were computed across the cortex and fitted using theoretical variograms to estimate sill and effective range parameters at each vertex. The resulting vertex-wise sill and range maps are shown in **Fig. 4B**. Consistent with previous findings (Leech et al., 2023), unimodal regions such as the visual cortex exhibited lower sill values and longer effective ranges, consistent with lower spatial variability and spatial dependencies extending over longer cortical distances. In contrast, association regions, particularly within the frontoparietal network, showed higher sill values and shorter effective ranges, indicative of greater local spatial variability and more spatially restricted spatial dependencies.

To examine the relationship between SWM fiber architecture and spatial autocorrelation properties, radial-tangential fiber predominance map was correlated with variogram metrics (**Fig. 4C**). This map was generated by subtracting the z-scored tangential AFD map from the z-scored radial AFD map at the 2 mm SWM depth, such that negative values indicated greater tangential fiber density and positive values indicated greater radial fiber density. Radial-tangential fiber predominance map showed significant positive correlations with both effective range (*r*=0.45, *P_eigen_*=0.001) and sill (*r*=0.32, *P_eigen_*=0.008) (see also **Supplementary Fig. 3**). These findings indicate that association cortices characterized by steeper variograms were associated with greater tangential fiber density, whereas unimodal cortices with more gradual variograms were associated with greater radial fiber density.

## DISCUSSION

In this study, we focused on separating radially and tangentially oriented fiber properties within the human SWM using *in-vivo* 3T and 7T MRI to characterize SWM network organization, as a proxy to dissociate long-range cortico-cortical connections and short-range U-fibers. We implemented a surface-based and reproducible framework for sampling SWM intensity profiles from multiple fiber-related features in the human brain while maintaining minimal dependencies to ensure broad accessibility. By dissociating fODF lobes relative to the Laplacian streamlines across different SWM depths, we generated AFD maps for radial and tangential components and subsequently derived FA and MD maps applying the MRDS framework. The validity of radial-tangential fiber component separation was supported by tractography-based bundle analyses, while the anatomical reliability of the tractography reconstructions used for these analyses was confirmed through comparison with *post-mortem* WM dissection data. To investigate the functional relevance of these fiber populations, we examined the relationship between spatial patterns of functional connectivity dissimilarity and microstructural predominance between radial and tangential fiber density. Notably, greater tangential fiber density was associated with regions exhibiting more rapid changes in functional similarity across cortical space, whereas greater radial fiber density was linked to more gradual functional transitions. This pattern may reflect the role of short-range U-fibers, which are oriented tangentially to the Laplacian streamlines and may support local connectivity between neighboring cortical regions with distinct functional profiles. Together, these findings underscore the complex organization of SWM and provide a foundation for investigating the distinct contribution of radial and tangential fiber component properties in neuroimaging studies, with potential implications for understanding how local white matter architecture and connectivity support the fine-scale functional organization that underlies large-scale brain networks and cognition.

The first 2 mm of the SWM compartment are characterized by high structural complexity, as demonstrated by our previous work showing pronounced depth-dependent microstructural variation within this superficial compartment (Hwang et al., 2026), which likely reflects the interplay between tangentially oriented U-fibers running parallel to the GM/WM interface and radially oriented long-range cortico-cortical afferents and efferents intersecting it perpendicularly (Van Dyken et al., 2024; Garcia-Saldivar et al., 2024). However, these previous measurements characterized the SWM as a composite compartment and did not distinguish the contributions of different fiber populations to its microstructural organization. In the present study, we extend this work by separating the fiber populations based on the assumption that U-fibers predominantly follow the GM/WM interface whereas long-range fibers approach the cortex more radially within the SWM. We therefore attributed each fODF lobe, representing a distinct fiber population within a voxel, according to its orientation relative to the Laplacian streamlines. Lobes aligned parallel to the Laplacian streamlines were assigned to the radial fiber component, whereas the remaining lobes were assigned to the tangential fiber component. Note that the radial component examined here was defined within the 3 mm depth of SWM and therefore may capture only the proximal portions of long-range projections rather than their full trajectories through the deep white matter. The observed effects should therefore be interpreted as reflecting the interaction between local SWM organization and the sampled components of extended cortico-cortical connections. Our results in **Fig. 2** revealed higher tangential fiber density in paralimbic and peri-central regions. Notably, paralimbic regions, which occupy an intermediate position between heteromodal association cortex and limbic areas within Mesulam’s cortical hierarchy (Mesulam et al., 1998), demonstrated relatively higher tangential AFD alongside lower radial AFD across SWM depths, suggesting a preferential increase in tangentially oriented superficial connectivity relative to radially organized fibers. Interestingly, radial AFD in **Fig. 2** showed a very similar pattern of intensity changes across SWM depths within Mesulam’s classes to that of the T1 map, as demonstrated in our previous work (Hwang et al., 2026), where this pattern was linked to myelination across SWM depth. This suggests that the depth-dependent variation observed in the T1 map may be substantially reflected in the radial fiber component. In parallel, the pronounced decrease in MD with increasing SWM depth in paralimbic regions may reflect greater microstructural restriction and tissue complexity, potentially arising from dense local fiber interactions and crossing configurations (Beaulieu, 2002, Hwang et al., 2026).

From a developmental perspective, cortical folding reflects the interplay between multiple mechanical and biological constraints, with axonal tension along cortico-cortical connections representing one potential contributor. Such forces may link short-range connectivity patterns to gyral geometry, whereas long-range fibers are governed by additional wiring constraints (Van Essen, 1997). Although axonal tension is unlikely to fully explain gyrification, this framework provided an early hypothesis for how wiring constraints may couple cortical folding with the organization of corticocortical connectivity. This principle is reflected in **Fig. 3A**, where radially and tangentially oriented fiber connections exhibit systematic angular reorganization as a function of proximity to the GM/WM interface, consistent with geometry-dependent fiber organization. Importantly, anatomical validation against *post-*mortem dissection provides an independent reference for evaluating the reliability of tractography-based reconstructions in this anatomically complex compartment. Because *in-*vivo tractography is inherently sensitive to factors such as crossing fibers, tract geometry, and reconstruction parameters, direct comparison with anatomically confirmed white matter pathways can help assess whether the reconstructed bundles capture plausible anatomical trajectories and spatial extents. In the present study, the high spatial correspondence between *in-*vivo tractography and *post-mortem* dissection annotations (**Fig. 3B**) supports the anatomical reliability of the representative fiber pathways used to validate the proposed radial-tangential orientation framework. However, given that SWM organization closely follows cortical folding geometry (Hwang et al., 2026; Reveley et al., 2015), the elevated tangential fiber component derived from fixel-based orientation decomposition may partly reflect the complex gyral-sulcal architecture underlying local cortico-cortical connectivity, rather than solely differences in large-scale cortical hierarchy. In particular, highly curved and deeply folded cortical territories, such geometry may preferentially enhance tangential fiber organization immediately beneath the GM/WM interface. Consistent with this interpretation, previous anatomical studies have demonstrated prominent U-fiber systems around the pre- and postcentral gyri (Shinohara et al., 2020), alongside substantial regional heterogeneity across the cortex. In the cingulate cortex, the cingulum bundle courses tangentially along the cortical mantle and therefore contributes to the tangential fiber component despite being classically described as a long-range association tract (**Fig. 4C**) (Jones et al., 2013; Bubb et al., 2018). Its highly coherent and densely packed structure may cause it to resemble a continuous bundle of closely aligned short association fibers, rather than a collection of discrete U-fibers. Importantly, tangential orientations in the SWM do not necessarily imply a uniform underlying fiber architecture. At the gyral crown and sulcal fundus, axonal populations may be intrinsically intermingled due to local geometric constraints, as also suggested by the gyral-sulcal analysis in **Fig. 3A**. For example, while projection long-range fiber bundles may exhibit clear orientation differences from U-fibers, association long-range fiber tracts may show more similar angular characteristics to U-fiber systems due to their trajectory along the cortical mantle and anatomical configuration. This highlights an inherent limitation in fully disentangling short- and long-range fiber populations using macro-scale diffusion-based angular decomposition approaches, which cannot fully separate these fiber systems in the presence of complex geometric and anatomical overlap.

Our understanding of SWM architecture was extended by linking microstructural asymmetries to functional variability across the cortex. Given its close anatomical proximity to the cortex and its role in supporting cortical-cortical connections, the SWM may provide a substrate for local cortical organization. Because large-scale functional organization exhibits strong spatial dependencies across the cortex (Margulies et al., 2016), assessing spatial autocorrelation offers a meaningful framework for testing whether SWM microstructural predominance relates to similar organizational principles (Huntenburg et al., 2018, Leech et al., 2023). Indeed, regions with greater radial fiber contributions showed larger spatial dependency ranges, indicating more gradual changes in functional similarity across geodesic distance. In contrast, areas with greater tangential fiber contributions exhibited higher sill and lower ranges, consistent with more rapid local functional differentiation (**Fig. 4B-C**). These patterns broadly align with a hierarchical cortical organization in which unimodal regions display gradual functional changes with brain space, while association cortices show sharper local transitions reflecting increased microstructural and functional heterogeneity (Paquola et al., 2025; Leech et al., 2023). However, this relationship was not uniform across systems. Transmodal regions such as the frontoparietal and default mode networks showed both elevated sill and appreciable radial fiber contributions, suggesting a more complex interplay between local and distributed integration (Buckner & Krienen, 2013; Sepulcre et al., 2010). This observation is consistent with evidence that association cortex supports a coexistence of dense local cortico-cortical interactions and distributed integrative pathways underlying flexible cognition (Buckner & Krienen, 2013; Mesulam et al., 1998). In this context, U-fibers may support the local connectivity underlying fine-scale functional organization, including the coexistence of neighboring cortical territories with distinct functional profiles, whereas long-range fibers facilitate integration across distributed cortical systems (Catani et al., 2012; Wang et al., 2023). Such an interpretation is consistent with evidence from primate neuroanatomy and human neuroimaging demonstrating that association cortex contains spatially interdigitated functional networks, in which neighboring cortical patches can exhibit distinct functional identities despite their close anatomical proximity (Goldman-Rakic, 1988; Braga & Buckner, 2017). However, SWM microstructure alone cannot fully account for the spatial organization of functional connectivity. Variogram-derived metrics likely reflect the combined influence of multiple organizational axes, including cytoarchitectonic differentiation, topographic constraints, and large-scale network hierarchy, in addition to SWM architecture (Huntenburg et al., 2018). For example, limbic and insular regions demonstrated strong tangential fiber component predominance alongside short functional dependency ranges, whereas sensorimotor cortex exhibited relatively smoother functional transitions despite substantial tangential fiber contributions (**Fig. 4C**). Together, these results suggest that SWM provides an important but incomplete substrate for cortical functional geometry, emphasizing the need to consider multi-scale structural determinants when interpreting structure-function relationships in the human brain.

These findings highlight the potential structural and functional relevance of tangential fiber components as a promising feature for advancing cognitive and systems neuroscience research. A central objective of this work was to promote accessibility and reproducibility in SWM research through an open-science framework. To this end, we shared an open-access pipeline for dissociating tangential fiber component from radial fiber populations within the structurally complex SWM, while leveraging publicly available datasets to facilitate transparency and broader community engagement. The proposed framework was further supported by demonstrating concordance with *post-mortem* anatomically confirmed *in-vivo* diffusion tractography. Moreover, integrating microstructural predominance measures with spatial patterns of functional variability provides new insight into the potential contribution of tangential fiber components to local cortical organization in the human brain. Collectively, these efforts establish a foundation for future open-science investigations of SWM organization and provide methodological tools that may support emerging applications in precision neuroscience and neuroimaging research.

## MATERIALS AND METHODS

### 1. MRI data acquisition and preprocessing

All *in-vivo* MRI data were acquired at the McConnell Brain Imaging Centre of the Montreal Neurological Institute using Siemens MRI systems. The cohort consisted of eleven healthy participants (4 males, 7 females) who underwent both 3T and 7T MRI scans, with a mean age of 28.5±5.43 (mean±SD). Participants were recruited between March 2022 and March 2025. All MRI data correspond to a subset of participants from the MICA-MICs 3T MRI (Royer et al., 2022) and MICA-PNI 7T MRI (Cabalo et al., 2025) datasets. Detailed acquisition parameters and preprocessing workflows are provided in the publicly available dataset documentation (MICA-MICs: https://osf.io/j532r/ and https://portal.conp.ca/dataset?id=projects/mica-mics; MICA-PNI: https://osf.io/mhq3f/).

The following scans were utilized for each MRI session:

#### a. *In-vivo* 7T MRI acquisition

Imaging at 7T was performed on a Siemens Magnetom Terra scanner equipped with a 32-channel receive and 8-channel transmit head coil operating in parallel transmission (pTX) mode. Structural images were acquired using a three-dimensional magnetization-prepared two rapid gradient-echo sequence (Marques et al., 2010; Van Damme et al., 2021) (MP2RAGE; 0.5 mm isovoxels, matrix = 320×320, 320 sagittal slices, TR = 5170 ms, TE = 2.44 ms, TI1 = 1000 ms, TI2 = 3200 ms, flip = 4°, iPAT = 3, partial Fourier = 6/8, FOV = 260×260 mm^2^). Resting-state functional MRI data were collected using a multi-echo 2D BOLD echo-planar imaging sequence (Uğurbil et al., 2013) (University of Minnesota, CMRR; 1.9 mm isovoxels, 75 slices oriented to AC-PC-39 degrees, TR = 1690 ms, TE = 10.80/27.3/43.8 ms, flip = 67°, FOV = 224×224 mm^2^, slice thickness = 1.9 mm, MB = 3, echo spacing = 0.53 ms). During the resting-state acquisition, participants instructed to fixate on a gray cross and remain at rest for 7 min 45 s, resulting in 275 volumes.

#### b. *In-vivo* 3T MRI acquisition

Diffusion-weighted MRI was collected at 3T on a Siemens Magnetom Prisma-Fit system using a 64-channel phased-array head coil, as temporal lobe signal drop-off at 7T limited the reliability of the corresponding 7T data in that region. Three distinct diffusion shells were acquired with b-values of 0, 300, 700, and 2,000 s/mm^2^, with 10, 40, and 90 diffusion directions per shell, respectively. In addition, one reverse phase-encoded (posterior-to-anterior) b = 0 image was acquired for correction of susceptibility-induced distortions. Imaging parameters included 1.6 mm isovoxels, TR = 3500 ms, TE = 64.40 ms, flip angle = 90°, refocusing flip angle = 180°, FOV = 224 x 224 mm^2^, slice thickness = 1.6 mm, multi-band factor = 3, and echo spacing = 0.76 ms. Additional analyses using the available 7T diffusion data are presented in **Supplementary Fig. 1**.

#### **c.** Multimodal MRI preprocessing

In this study, we analyzed 7T structural and functional MRI from MICA-PNI (Cabalo et al., 2025) and 3T diffusion-weighted MRI from MICA-MICs (Royer et al., 2022). (1) Structural T1-weighted (T1w) images derived from MP2RAGE (Marques et al., 2010) were first deobliqued and reoriented to the standard LPI orientation (left-to-right, posterior-to-anterior, inferior-to-superior). The structural images were subsequently registered to both volumetric and surface-based templates. Geodesic distances between all cortical vertices on each subject’s native fsLR-5k mid-surface were then computed using Dijkstra’s algorithm (Dijkstra et al., 2022). (2) Resting-state functional MRI data were also reoriented to LPI orientation. For multi-echo acquisitions, time series were extracted from all echoes, nuisance signals were removed, and echoes were optimally combined. The resulting data were then decomposed into components using principal and independent component analysis with TEDANA v.0.0.12 (DuPre et al., 2021). Global signal regression was then performed to regress residual global fluctuations before functional connectivity estimation (Power et al., 2018, Cruces et al., 2022). Preprocessed time series underwent high-pass temporal filtering and were mapped onto the native cortical surface. Functional connectivity matrices were subsequently computed for each participant by cross correlating the time series across all nodes. Correlation coefficients were transformed using Fisher’s r-to-z transformation, and functional cortico-cortical connectivity matrices were generated based on the fsLR-5k symmetric surface template through subject-specific resampling.

(3) Diffusion-weighted MRI data were preprocessed in native space using *MRtrix3* (https://mrtrix.readthedocs.io/en/latest/) (Tournier et al., 2019), including Marchenko-Pasteur principal component analysis (MP-PCA) denoising (Cordero-Grande et al., 2019; Veraart et al., 2016), Gibbs ringing correction (Kellner et al., 2016; Andersson et al., 2003), head motion and susceptibility distortion correction, eddy current correction, and non-uniformity bias field correction (Andersson et al., 2003, 2016; Tustison et al., 2010; Smith et al., 2004). Fiber orientation distributions (FODs) were estimated using the multi-shell multi-tissue constrained spherical deconvolution (MSMT-CSD) algorithm (Jeurissen et al., 2014). To enable quantitative comparisons within the SWM, normalized FODs were generated using averaged tissue-specific response functions.

All MRI images were aligned to native *FastSurfer* (Henschel et al., 2020) space of each participant using label-based affine registration. Multimodal MRI pre- and post-processing were conducted using the open-access *micapipe* toolbox (version 0.2.3) (http://micapipe.readthedocs.io) (Cruces et al., 2022).

### 2. Separation of radial and tangential fiber components in the superficial white matter

To investigate the SWM, we applied an equipotential-based sampling framework designed to improve spatial smoothness and practical implementation while reducing artifacts and minimizing the need for extensive *post-hoc* corrections (Liu et al., 2016; Hong et al., 2019; Hwang et al., 2026). Specifically, the Laplace equation was solved within the white matter domain by first generating a Laplacian field between the GM/WM interface and the ventricular walls. An existing GM/WM surface was then shifted along the resulting gradient field, with termination criteria defined by the geodesic distance traveled. The resulting potential field allowed the systematic definition of SWM depths beneath the GM/WM interface. The code used to generate the Laplacian field using T1w image is publicly available in our open repository (DeKraker et al., 2024; Hwang et al., 2025). Based on this Laplacian field, vertex-wise streamlines were generated from the fsLR-32k GM/WM surface toward subcortical regions using 200 integration steps (step size = 0.1 mm) for streamline tracing. To transform the streamlines into diffusion MRI space, the MP2RAGE-T1w image was first nonlinearly registered to the b0 image using Advanced Normalization Tools (*ANTs*) (Avants et al., 2008), and the resulting inverse deformation field was applied to the streamlines using MRtrix3 (Tournier et al., 2019) (**Fig. 1A**). The streamlines were resampled with a 0.5 mm step size and subsequently truncated by 5 mm. These streamlines are termed Laplacian streamlines in this study to distinguish them from tractography-derived streamlines used in subsequent analyses.

The maximum SWM depth of 3 mm was chosen to capture both the U-fiber system and the termination zones of long-range bundles (Schüz et al., 2002, Hwang et al., 2025). To dissociate and quantify radial and tangential fiber components within the first 3mm of the SWM, we performed a diffusion fixel-based analysis using a MSMT-CSD model (Jeurissen et al., 2014). Fiber populations were classified based on their orientation relative to the Laplacian streamlines (Garcia-Saldivar et al., 2024). For each streamline segment, the dot product between the segment direction and the orientations of the underlying fODF lobes, representing distinct fiber orientations within a voxel, was computed. The fODF lobe with the largest dot product with the streamline direction and angular deviations of less than 40° in gyral vertices and 50° in sulcal vertices were classified as the parallel component (see **Figure 3A**). The remaining fODF lobes were averaged to represent the perpendicular component. Based on this classification, the parallel component was interpreted as radial oriented fibers projecting afferently and efferently to the cortex from deeper white matter (*i.e.*, long-range fibers), whereas the perpendicular component was considered to represent fibers oriented tangentially to the GM/WM interface, consistent with putative U-fibers. AFD (Raffelt et al., 2012) was then quantified as the integral of each fiber population’s FOD lobe across SWM depths (**Fig. 1B**). The resulting AFD maps were subsequently downsampled to 5k vertices per hemisphere using *connectome* workbench (Marcus et al., 2013) to enable functional analyses. All code implementing this pipeline is openly available at https://github.com/lconcha/corticalDWI.

### 3. Bundle-specific diffusion metric mapping using multi-tensor

In addition to fixel-based density estimates, we applied a MRDS approach (Coronado-Leija et al., 2017) to derive diffusion tensor metrics for radial and tangential fiber components separately. The MRDS strategy enables the estimation of distinct diffusion profiles for individual fiber bundles within a voxel through a hierarchical multi-resolution search over a discrete orientation set, allowing accurate and computationally efficient orientation estimation. Following this strategy, between one and three diffusion tensors were estimated per voxel using an F-test model to characterize bundle-specific diffusion profiles in nested models. Using the orientation criteria described above, the resulting tensors were then classified into the tensor most parallel to the Laplacian streamline direction and the average of the remaining tensors, corresponding to radially and tangentially oriented fiber populations, respectively. Diffusion metrics, including FA and MD, were then computed independently for each diffusion tensor and sampled across multiple SWM depths to characterize depth-dependent microstructural differences within the SWM.

### 4. SWM contextualization with cortical organization

SWM intensity profiles derived from multiple diffusion metrics were interpreted in relation to normative patterns of cortical organization, reflecting regional differences in laminar differentiation and hierarchical specialization across the cortex. Profiles were grouped according to cortical classes defined by Mesulam atlas (Mesulam et al., 2000), which categorizes cortical regions into four types based on the degree of laminar differentiation, hierarchical organization, and functional specialization. To relate SWM features to cortical organization, Laplacian field mapping was used to establish an isomorphic correspondence between the cortical surface and adjacent SWM, enabling cortical labels to be propagated from surface vertices into the underlying white matter.

### 5. Anatomical assessment using diffusion MRI tractography and *post-mortem* dissection photogrammetry

#### a. Subject-specific tractography

For the validation analysis, we first sought to determine whether anatomically plausible radial and tangential fiber systems within the SWM could be distinguished based on their orientations relative to the Laplacian streamlines. To this end, tractography reconstruction was focused on the connectivity between PrCG and PoCG as a representative tangentially oriented fiber bundle, and between the MFG and AG as a representative radially oriented fiber bundle. These connections were selected to correspond to manually annotated white matter pathways from *post-mortem* brain dissection photogrammetry dataset (Vavassori et al., 2025b), allowing the assessment of the anatomical reliability of the selected representative white matter bundles through direct comparison between *in-vivo* tractography and *post-mortem* dissection.

The Desikan-Killiany atlas (Desikan et al., 2006), generated during the segmentation step using FastSurfer (Henschel et al. 2020), was used for cortical and white matter parcellation. Using the resulting gray matter and white matter parcellation, probabilistic tractography was performed with the iFOD2 algorithm implemented in MRtrix3 (Tournier et al., 2019) to investigate the structural connectivity between the PrCG and PoCG, and between the MFG and AG. Tracking parameters for the PrCG-PoCG connectivity were as follows: minimum/maximum streamline length = 30/60 mm, step size = 0.5 mm, maximum curvature angle = 60°, and FOD cutoff for streamline termination = 0.1. Streamlines were seeded from the precentral gyrus white matter and constrained to pass through both the precentral and postcentral gyri while remaining within the first 3 mm of the SWM. For MFG-AG connectivity, streamlines were seeded from the inferior parietal lobule white matter and required to connect the middle frontal gyrus and inferior parietal lobule while excluding the temporal lobe, cingulate cortex, insula and subcortical structures. Tracking parameters were: minimum/maximum streamline length = 60/180 mm, step size = 0.5 mm, maximum curvature angle = 45°, and FOD cutoff = 0.1. For both pathways, 10,000 streamlines were generated per subject. Complete tractography reconstruction criteria are provided in **Supplementary Table 3**.

Streamline seeding was performed within the white matter, as diffusion-weighted signal intensity in the cortical gray matter is relatively low and may lead to distortions at the GM/WM interface. In addition, streamline endpoints crossing the GM/WM boundary were precisely cropped, and anatomically constrained tractography was applied to ensure anatomically plausible streamlines terminating within the gray matter (Smith et al., 2013). Optimal step size and maximum curvature angle were determined through a parameter optimization procedure based on streamline reconstruction performance across different seed densities. Subject-specific tractography bundles derived from 11 subjects were combined into a single population template to enable anatomical evaluation using brain dissection photogrammetry.

#### **b.** Anatomical evaluation with *post-mortem* dissection photogrammetry

Anatomical evaluation of tractography templates was performed following the integrated *in-vivo* tractography and *post-mortem* dissection anatomical validation pipeline (Vavassori et al., 2025a), based on *post-mortem* brain dissection photogrammetry data from the *BraDiPho* repository (https://bradipho.eu). The dataset included ten multi-layered photogrammetry models of *post-mortem* dissection (6 left-hemisphere and 4 right-hemisphere) performed according to Klingler’s method and registered to the specimen T1w MRI, enabling direct comparison between tractography-derived templates and manually annotated white matter pathways across sequential dissection layers. Tractography templates were non-linearly transformed into the *post-mortem* specimen space using *ANTs* (Avants et al., 2009) (left-hemisphere templates to left-hemisphere specimens and right-hemisphere templates to right-hemisphere specimens) and rendered within the photogrammetric 3D models for conjoint evaluation with anatomical annotations derived from dissection in *CloudCompare* (version 2.13.2) (http://www.cloudcompare.org/).

Qualitative assessment was first performed by visually comparing the anatomical trajectories and spatial extent of tractography bundles with manually annotated white matter pathways across successive dissection layers. Because the tractography templates were derived from a population-based dataset whereas the dissection data originated from individual specimens, the evaluation focused on not only on confirming the existence of specific cortical connections defined by their cortical terminations, but also on assessing whether the anatomical extent of the dissected pathways was encompassed by the tractography reconstruction.

To complement qualitative inspection, quantitative spatial overlap analysis was conducted using signed-distance computations implemented in *CloudCompare*. First, distances between tractography templates and dissection surfaces were computed to identify regions of close spatial correspondence, and points located within 1 mm of the dissection surface were retained to generate an overlap mesh between tractography and dissection representations. A second signed-distance computation was subsequently performed between this overlap mesh and the manual annotation of the corresponding connection in each dissection model fiber pathways to quantify local discrepancies between the two modalities. Agreement was then estimated as the proportion of annotation points in the annotation mask located within 1 mm of the tractography-defined overlap region. This procedure enabled assessment of the anatomical reliability of the tractography templates representing radial and tangential white matter connections, which were subsequently used for further validation analyses.

#### **c.** Fiber orientations relative to Laplacian streamlines

To quantitatively validate the orientation classification framework, we assessed whether tractography-defined radial and tangential fiber bundles exhibited the distinct angular relationships relative to the Laplacian streamlines within the SWM. Streamline orientations from anatomically defined tractography bundles were compared with local Laplacian streamline directions within the first 3 mm of the white matter. The anatomically evaluated representative fiber pathways described above were included in this analysis. Analyses were performed independently for the left and right hemispheres using an identical procedure.

For each subject, tractography streamlines were compared with subject-specific Laplacian streamlines. Local Laplacian orientations were extracted separately within three SWM depth intervals (0-1 mm, 1-2 mm, and 2-3 mm from the GM/WM interface). Depth along each Laplacian streamline was estimated using cumulative Euclidean distance from the streamline origin at the GM/WM interface.

For each tractography sampling point, the local streamline orientation was compared with the nearest Laplacian streamline direction within the corresponding SWM depth interval. Only streamline points located within 1 mm of a Laplacian reference point were retained for angular analysis. Angular similarity between tractography streamlines and Laplacian streamlines was quantified using the absolute dot product between normalized tangent vectors. Mean angular values were computed separately for each SWM depth interval and subject.

### 6. Empirical and theoretical variograms construction

Cortical functional organization exhibits spatial dependencies, with functional similarity between cortical regions varying systematically as a function of anatomical distance (Leech et al., 2023). Given that SWM fiber populations support both local cortico-cortical interactions and long-range communication, these spatial dependencies provide a framework for assessing how SWM architecture is reflected in functional organization. We specifically calculated empirical variograms (Leech et al., 2023), which quantifies how rapidly functional organization changes with spatial distance. Pairwise geodesic distance between cortical vertices were grouped into 20 uniformly spaced distance bins, with the maximum distance restricted to 160 mm (all vertex pairs separated by distances up to 160 mm were included in the analysis – see **Supplementary Fig. 4**). Functional dissimilarity was defined as (1 - *Pearson’s correlation coefficient*) computed between vertex-wise functional connectivity matrix, forming a functional dissimilarity matrix. These values were subsequently aggregated within distance bins using Gaussian smoothing (Burt et al., 2020; Viladomat et al., 2014), yielding a whole-brain empirical variogram (**Fig. 4A**). For vertex-wise variograms, the same procedure was applied independently to each row of the functional connectivity and geodesic distance matrices, producing a localized variogram estimate per vertex. The empirical variogram captures the rate at which functional similarity varies with geodesic distance across the cortical surface, both globally and locally. Whole-brain and vertex-wise variograms were computed using openly available code (https://github.com/ActiveNeuroImaging/BrainVariograms) (Leech et al., 2023).

Theoretical variograms were fitted to the empirical variograms to provide a compact summary of the spatial autocorrelation structure using a small number of parameters, thereby facilitating comparisons across networks and hemispheres. Based on prior studies demonstrating exponential relationships between spatial distance and neural measures (Leech et al., 2023; Shinn et al., 2023), empirical variograms were characterized using an exponential model (**Fig. 4A, bottom right**). Specifically, the empirical variograms were fitted using a non-linear least squares approach to estimate the sill and range parameters of the exponential. The sill represented the asymptotic level of functional dissimilarity, and the effective range represented the spatial distance at which the variogram reached 95% of the sill value. To minimize the influence of sparsely sampled distances and local noise effect, empirical variograms were trimmed to bins between 2 and 19 prior to model fitting. Statistical significance was evaluated using eigenstrapping-based surrogate maps (1,000 permutations) to account for spatial autocorrelation (Koussis et al., 2025).

## DATA AND CODE AVAILABILITY

The 3T and 7T data are available on the Open Science Framework at https://osf.io/j532r/ and https://osf.io/mhq3f/. The code implementing the processing pipeline is openly available on GitHub at https://github.com/lconcha/corticalDWI, with a tutorial at https://github.com/lconcha/corticalDWI/blob/main/tutorial.md. The code and Jupyter notebook for reproducing the analyses are available at https://github.com/y0ungeunii/radial-tangential_fiber_systems_in_swm.

