## Supplementary for "RADIAL AND TANGENTIAL FIBER COMPONENTS IN HUMAN SUPERFICIAL WHITE MATTER AND THEIR ASSOCIATIONS WITH MACROSCALE FUNCTION"

**Supplementary materials**

**Dissociating U-fibers from long-range fibers
in human superficial white matter
and their associations with functional variability**

Youngeun Hwang^1,2^, Laura Vavassori^1,2^, Paul Bautin^1,2^, Raul Rodriguez-Cruces^1,2^, Edith Gaspar-Martinez^3^, Donna Gift Cabalo^1,2^, Alexander Ngo^1,2^, Meaghan Smith^1,2^, Robert Leech^4^, Ricardo Coronado-Leija^5^, Alan C Evans^2^, Luis Concha^3^, Boris C. Bernhardt^1,2^

**Author affiliations:**

*^1^Multimodal Imaging and Connectome Analysis (MICA) Lab and Centre for Excellence in Epilepsy at the Neuro (CEEN), McGill University, Montreal, QC, Canada,*

*^2^McConnell Brain Imaging Centre, Montreal Neurological Institute and Hospital, Montreal, QC, Canada,*

*^3^Institute of Neurobiology, Universidad Nacional Autónoma de México (UNAM), Querétaro, Querétaro, Mexico,*

*^4^King’s College London, London, UK,*

*^5^Center of Research in Mathematics, Guanajuato, Mexico*

**Corresponding authors:**

- Youngeun Hwang, MSc
- Boris C. Bernhardt, PhD

**Supplementary tables and figures**

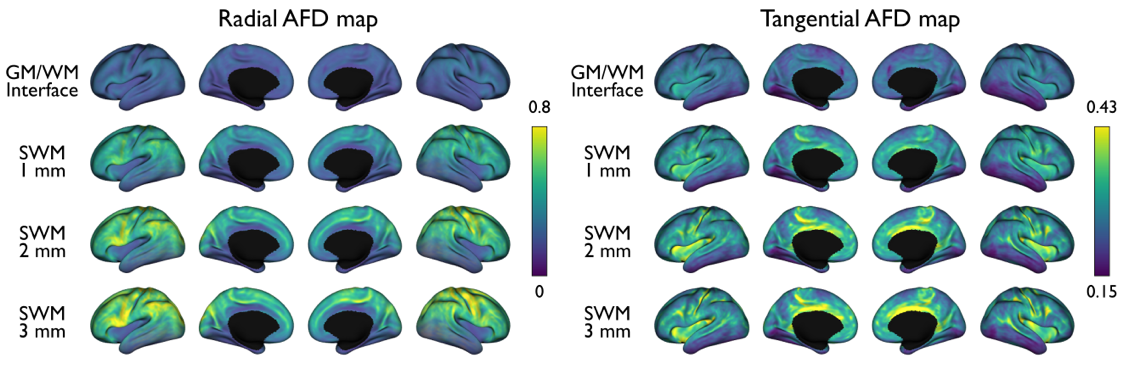

**Supplementary Fig. 1.** Radial and tangential AFD maps derived from the available 7T diffusion MRI data. The same analysis was applied as for the 3T data. However, temporal lobe signal drop-off at 7T limited the reliability of the corresponding results.

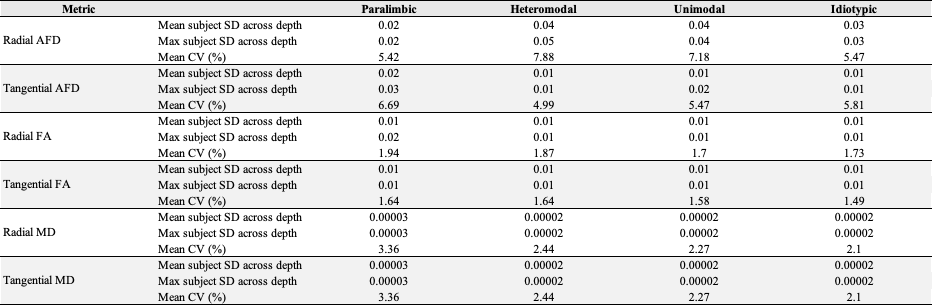

**Supplementary Table 1.** Subject-level variability was quantified using the coefficient of variation (CV), defined as the ratio of the standard deviation (SD) across subjects to the mean profile. CV was computed for radial and tangential AFD, FA, and MD profiles across SWM depths (0-3mm), within each cortical class.

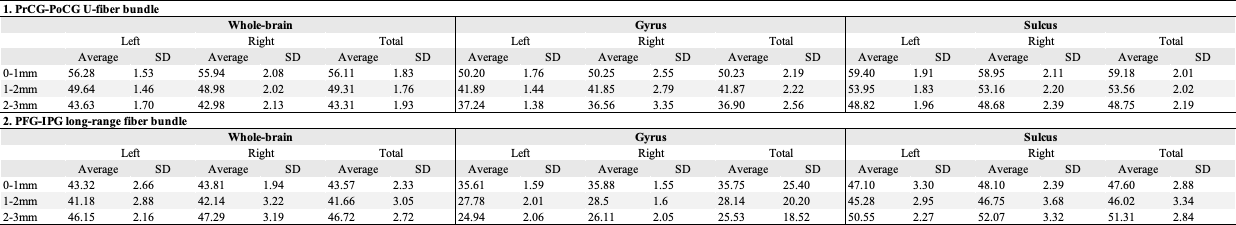

**Supplementary Table 2.** Angles between representative short- and long-range fiber tracts and Laplacian streamlines were computed at gyral and sulcal vertices across SWM depths (0-1 mm, 1-2 mm and 2-3 mm).

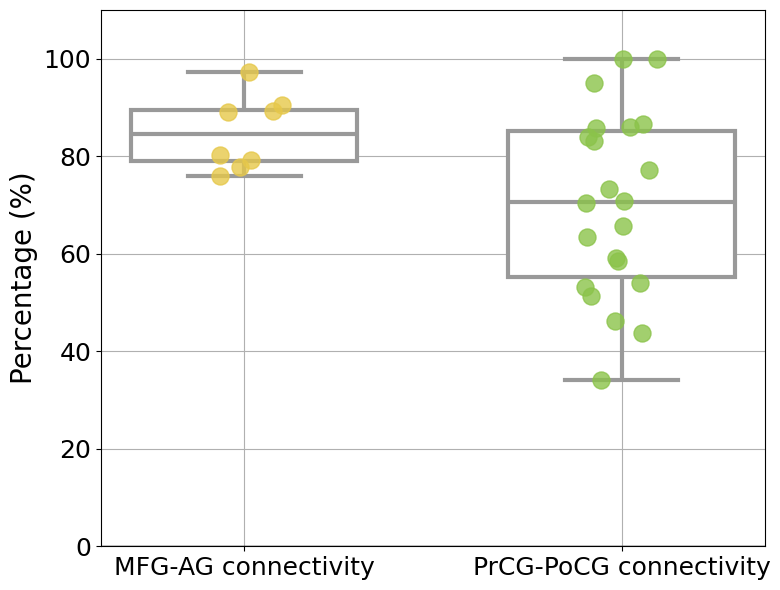

**Supplementary Fig. 2.** Percentage of overlap between tractography and corresponding annotation along different dissection epochs. This proportion considers the number of points of the manual annotation of the bundle that lay within a 1 mm distance from the mesh representing the overlap between the tractography reconstruction and the dissection epoch. Epochs showing the highest agreement between tractography and annotation are highlighted in red. **Abbreviations:** AF, angular gyrus; MFG, middle frontal gyrus, PoCG: postcentral gyrus; PrCG, precentral gyrus; spc, specimen; epo, epoch (layer of dissection).

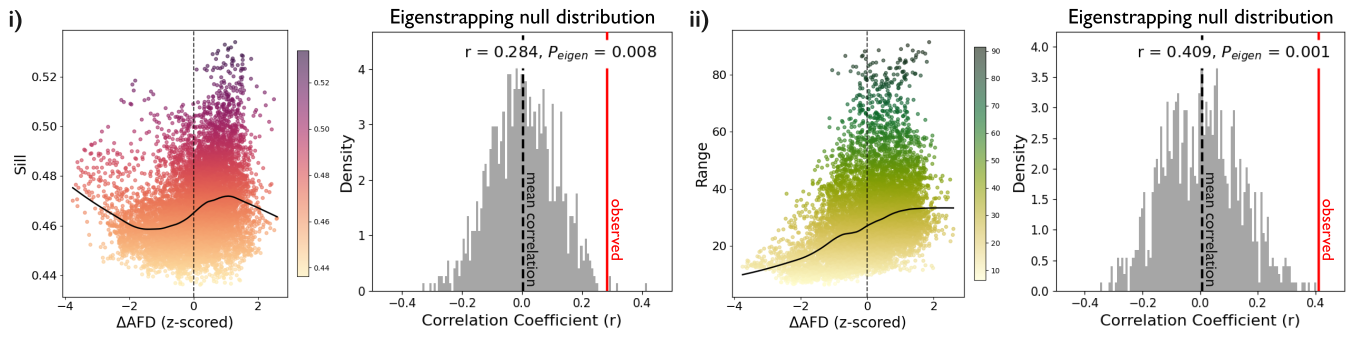

**Supplementary Fig. 3.** Correlations between variogram-derived metrics and microstructural asymmetry measures, with statistical significance assessed using an eigenstrapping permutation testing.

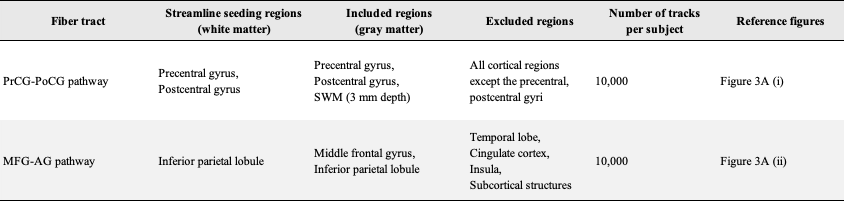

**Supplementary Table 3.** Tractography reconstruction criteria for PrCG-PoCG and MFG-AG pathways. **Abbreviations:** PrCG, precentral; PoCG, postcentral; MFG, middle frontal gyrus; AG, angular gyrus.

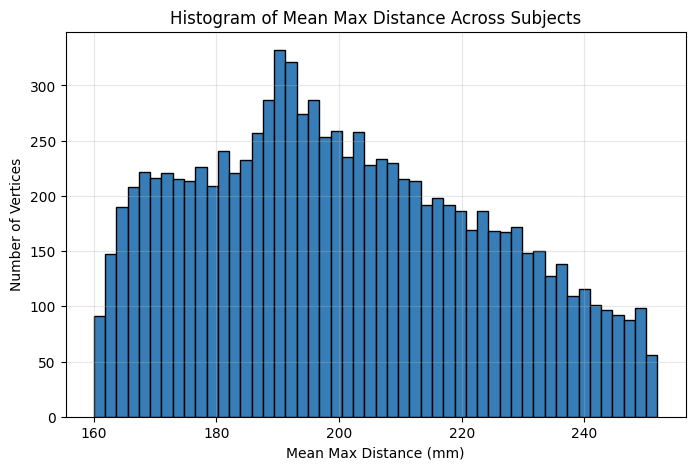

**Supplementary Fig. 4.** Distribution of maximum distances for each vertex.
